# Expanded adaptive NKG2C+ NK cells exhibit potent ADCC and functional responses against HBV-infected hepatoma cell lines

**DOI:** 10.1101/2025.05.29.655595

**Authors:** Jonida Kokiçi, Helena Arellano-Ballestero, Benjamin Hammond, Anucha Preechanukul, Noshin Hussain, Kelly da Costa, Jessica Davies, Sadiyah Mukhtar, Thomas Fernandez, Sabine Kinloch, Fiona M Burns, Patrick Kennedy, Douglas MacDonald, Mala K Maini, Upkar S Gill, Alan Xiaodong Zhuang, Mark W Lowdell, Karl-Johan Malmberg, Ebba Sohlberg, Dimitra Peppa

## Abstract

**Background:** Hepatitis B virus (HBV) infection remains a significant global health challenge, leading to chronic liver disease and hepatocellular carcinoma (HCC). Natural killer (NK) cells play an important role in the clearance of HBV-infected cells, but their efficacy is often compromised during chronic infection. Adaptive NK cells, characterised by NKG2C expression and enhanced functional responses, represent a promising therapeutic avenue for enhancing anti-HBV immunity and responses to HBV-driven cancers.

**Methods:** We applied an established protocol, involving K562-HLA-E expressing feeder cells and cytokines (IL-2), for the expansion of adaptive NK cells from cryopreserved T- and B cell depleted peripheral blood mononuclear cells (PBMCs) derived from donors with chronic HBV infection alone or with Human Immunodeficiency Virus (HIV) co-infection. We evaluated the adaptive profile of expanded NK cells, their antibody-dependent cellular cytotoxicity (ADCC) capacity and functional responses against hepatoma cell lines in the presence or absence of HBV infection.

**Results:** Expanded NK cells achieved >97% purity, with the NKG2C positive population exhibiting a mean 100-fold expansion. These cells demonstrated a predominantly adaptive phenotype with high surface expression of NKG2C and cytotoxic potential (Granzyme B). They maintained high levels of CD16 surface expression and upregulated CD2, essential for ADCC. Functionally, expanded adaptive NK cells showed enhanced ADCC capacity and functional responses to K562 targets, naive, HBV integrant-expressing, and *de novo* infected hepatoma cell lines. TGF-β preconditioning induced tissue-resident features (CD103, CD49a) in expanded adaptive NK cells, while preserving their adaptive phenotype and functionality, enhancing their potential for liver targeted immunotherapy. Further, expanded adaptive NK cells demonstrated minimal reactivity against autologous activated T cells, suggesting limited off-target effects.

**Conclusions:** Our study demonstrates the first successful expansion of adaptive NK cells with robust functional responses from donors with chronic viral infection. This approach creates opportunities for NK cell-based therapies alone or in combination with monoclonal antibodies contributing to HBV functional cure strategies and the treatment of HBV-driven cancers.

## Introduction

Hepatitis B virus (HBV) infection poses a significant global health threat, leading to substantial liver-related morbidity, accounting for more than 800,000 deaths per year globally [1]. Chronic HBV infection is a major contributor to hepatocellular carcinoma (HCC), which is not only the most prevalent type of liver cancer but also the third leading cause of cancer-related deaths worldwide [2, 3]. There is therefore an unmet need to improve current therapies to increase ‘functional’ cure for HBV and related co-morbidities.

Natural Killer (NK) cells are particularly concentrated within the liver and serve as a cornerstone of the immune defence, equipped to recognise and eliminate virus-infected and malignant cells [4, 5]. Studies on human and animal models have previously established the dual nature of NK cells in HBV infection, where they can both contribute to viral control and paradoxically to disease pathogenesis [6-10]. In chronic HBV, we have demonstrated that while conventional NK cells show compromised Interferon gamma (IFNγ) production, they maintain high expression of Tumour necrosis factor (TNF)-related apoptosis-inducing ligand (TRAIL) that can both directly mediate hepatocyte death and impair HBV-specific T cell responses in the liver [7, 8]. This functional dichotomy underscores the importance of identifying and harnessing distinct NK cell subpopulations with beneficial properties while minimising pathogenic effects to better orchestrate overall immune responses against HBV.

Of particular interest is the identification of NK cells with memory traits (so-called memory or adaptive NK cells) which exhibit marked phenotypic and functional parallels with cytotoxic CD8 T cells [11], suggesting significant potential for clinical exploitation over conventional NK cells [12]. These adaptive NK cells, most extensively characterised in the context of human cytomegalovirus (HCMV) infection, are defined by higher expression of NKG2C (an activating receptor that recognises HLA-E). They show enhanced functional responses, including increased antibody-dependent cellular cytotoxicity (ADCC) [13], and IFNγ production, attributed to epigenetic modification of the *IFNG* locus [11], with the ability to activate endogenous immune responses. Furthermore, adaptive NK cells possess single self-killer immunoglobulin-like receptors (KIRs), which can be leveraged to maximise ‘missing-self’ reactivity against tumours [14, 15]. They have also shown resilience against suppression by myeloid-derived suppressor cells (MDSC), which often impede effective NK cell reactivity [16]. Adaptive NK cells exhibit an elevated bioenergetic profile, with heightened glycolysis and OXPHOS compared to conventional NK cells, allowing them to better withstand the harsh microenvironment of inflamed tissues [17]. Furthermore, they also exhibit reduced immunoregulatory capacity, showing diminished degranulation when encountering activated T cells and limited capability to suppress virus-specific T cell responses [11, 18].

We have recently identified that adaptive NK cell subsets expressing NKG2C are prominent in people with HBV/HIV co-infection and potentially advantageous in controlling HBV [19]. These populations were characterised by superior functional responses compared to conventional NK cells and inversely correlated with circulating HBV-RNA and HBsAg levels, implying a vital role in HBV control [19]. This suggests that adaptive NK cells may be effective at targeting HBV-infected hepatocytes while potentially avoiding the detrimental effects mediated by conventional NK cells.

Building on these insights, we hypothesised that expanding adaptive NK cells from donors with chronic viral infection could generate populations with enhanced antiviral activity against HBV-infected cells. In this study, we applied a Good Manufacturing Practice (GMP)-compliant expansion protocol leveraging an HLA-E expressing K562 feeder cell line, to generate adaptive NK cells from donors with chronic viral infection. We assessed their phenotype and functional capacity and evaluated their potential as an innovative immunotherapeutic approach for HBV infection and HCC using *in vitro* models.

## Methods

### Human Participants and Cell Processing

A total of 30 patients with chronic viral infection (HBV or HBV/HIV), all suppressed on antiviral therapy with an undetectable HBV DNA and HIV viral load, were recruited at Mortimer Market Centre for Sexual Health and HIV research, the Ian Charleson Day Centre at the Royal Free Hospital (London, UK) or The Royal London Hospital (London, UK) following written informed consent as part of a study approved by the local ethics committee (Berkshire [REC 16/SC/0265] and London Bridge [REC 17/LO/0266]) and conformed to the Helsinki Declaration principles. All participants were confirmed HCV negative and HCMV seropositive.

Peripheral blood mononuclear cells (PBMCs), plasma, and serum were collected. Whole blood was collected in Heparin coated tubes, centrifuged for 5 minutes at 800g to collect plasma which was stored in aliquots at -80°C. The remaining volume of blood was diluted with Roswell Park Memorial Institute medium (RPMI) (Gibco, Paisley, UK) and then layered onto Histopaque-1077 gradient (Sigma Aldrich UK), centrifuged for 20 minutes at 800g without a brake. The resulting buffy coats containing PBMCs were then collected, washed with RPMI and centrifuged. Concentration of live PBMCs was ascertained using Trypan blue and counted using an automated cell counter (Biorad, Hercules, CA, USA). PBMCs were resuspended in freezing media (10% dimethyl sulfoxide (DMSO) (Honeywell, Seetze, Germany) and 90% heat inactivated foetal bovine serum (FBS) and cryopreserved at -80°C in a Mr Frosty container overnight before transfer to liquid nitrogen storage.

### Expansion Protocol for Adaptive NK Cells

K562-HLA-E cells were irradiated with a dose of 100Gy on a CellRad X-Ray Irradiator. Irradiated K562-HLA-E feeder cells were resuspended in serum-free Opti-MEM media (Gibco, Paisley, UK) and pulsed with 300uM G-leader peptide overnight. HLA-E expression was confirmed the following day via flow cytometry (clone 2D12, Biolegend).

Cryopreserved PBMCs were thawed, centrifuged and rested for 1 hour at 37°C in complete RPMI medium (RPMI supplemented with Penicillin-Streptomycin, L-Glutamine, HEPES, essential amino acids, non-essential amino acids, and 10% FBS. After resting, cells were centrifuged at 300g for 10 min at 4°C and resuspended in MACS buffer (PBS supplemented with 0.5% bovine serum albumin and 2mM EDTA) for subsequent CD3/CD19 depletion, using CD3 Microbeads (Miltenyi Biotec) and CD19 Microbeads (Miltenyi Biotec) as per manufacturer instructions using MidiMACS Separator (Miltenyi Biotec), LD columns (Miltenyi Biotec) and pre-separation filters (Miltenyi Biotec). Depleted PBMCs were resuspended in GMP-grade Stem Cell Growth Medium (SCGM) medium (Sartorius CellGenix) supplemented with 10% human AB serum (Merck) and 1% L-Glutamine (Sigma Aldrich). A small fraction of pre-depletion and post-depletion PBMCs were stained for extracellular markers (CD3, CD19, CD56, CD16, CD14) to confirm depletion and purity.

Depleted PBMCs were co-cultured with irradiated K562-HLA-E feeder cells at a 1:2 ratio in 24-well G-Rex plates (Wilson Wolf) at 0.5×10^6^ total cells/cm^2^ and supplemented with 100IU/ml recombinant human IL-2 (PeproTech) for 11 days with 60% medium change on day 7, and IL-2 addition on days 4, 7 and 10. In the TGF-β condition, cells were supplemented with 10ng/ml recombinant human TGF-β (PeproTech) on day 0, 4, 7 and 10. Expanded NK cells on day 11 were resuspended in complete RPMI medium and used as indicated.

### NK cell isolation

Cryopreserved PBMCs were thawed and rested for 1 hour at 37°C in complete RPMI medium.After resting, cells were centrifuged at 300g for 10min at 4°C and resuspended in MACS buffer (PBS supplemented with 0.5% bovine serum albumin and 2mM EDTA) for subsequent NK cell isolation, using NK isolation kit (Miltenyi Biotec) as per manufacturer instructions using MidiMACS Separator (Miltenyi Biotec), LS columns (Miltenyi Biotec) and pre-separation filters (Miltenyi Biotec). Untouched NK cells were resuspended in complete RPMI medium and used as indicated.

### Phenotypic assessment

*Ex vivo* day 0 and expanded day 11 NK cells underwent phenotypic assessment. Cells were washed with PBS and stained with fluorochrome conjugated antibodies in combinations as listed in Table S1 with live/dead dye (Invitrogen, UK) for 20 minutes at 4°C. For subsequent intracellular cytokine staining, cells were then fixed and permeabilized with Cytoperm/Cytofix (Becton Dickenson, Wokingham, UK) before staining with fluorochrome conjugated antibodies (Supplemental Table S1) for 30 minutes at 4°C. For intranuclear staining, cells were fixed and permeabilized with Foxp3/Transcription Factor Staining (eBioscience) diluted as per manufacturer’s instructions, before staining with anti-PLZF PE/CF594, anti-Granzyme B Alexa-fluor 700 and anti-FcεRIγ FITC fluorochrome conjugated antibodies for 45 minutes at room temperature. Samples were acquired using BD Fortessa x20 using BD FACSDiva 8.0 or Cytek Aurora, using Cytek SpectroFlo software and analysed on FlowJo10.10.0. The gating strategy used for flow cytometry analysis is provided in Supplemental Figure 1b. viSNE analysis was performed using Cytobank (https://www.cytobank.org).

### Functional Assays

Matched *ex vivo* NK cells versus expanded NK cells were assessed for ADCC and functional responses against K562 and hepatoma cell lines HepG2-NTCP and PLC/PRF/5, as well as HBV-infected HepG2-NTCP cells and HepG2.2.15 cells.

For ADCC assays, Raji cells expressing CD20 were incubated for 30 minutes with anti-CD20 antibodies (Invivogen, Ca, USA) or isotype control at final concentration of 5ug/ml. *Ex vivo* or expanded NK cells were co-cultured with antibody-coated Raji or isotype-coated Raji cells at an effector: target (E:T) ratio of 10:1. An E:T ratio of 5:1 was used for K562 cells, PLC/PRF/5, HepG2-NTCP and HepG2.2.15 cells. One well of unstimulated effector cells was also included for each donor. Anti-CD107a APC-H7 fluorochrome was added to each well at the time of stimulation. Following 1 hour incubation at 37°C, Golgiplug / Golgistop were added to each well and incubated for a further 5 hours. At the end of the incubation, 20μl supernatant was collected from each well for Toxilight readings for the indicated experiments. Cells were stained with fluorochrome conjugated antibodies (listed in Supplemental Table S1) for 20 minutes at 4°C. Cells were washed with PBS before being fixed for 20 minutes at 4°C with Cytofix (Becton Dickenson, Wokingham, UK) and permeabilised with Cytoperm (Becton Dickenson, Wokingham, UK). Intracellular Cytokine Staining (ICS) was carried out with anti-IFNγ BV421-conjugated antibody and anti-TNFα BV711-conjugated antibody diluted in Cytoperm buffer by incubating cells for 30 minutes at 4°C. Samples were acquired using a BD Fortessa x20 using BD FACSDiva 8.0 and analysed using FlowJo10.10.0. The gating strategy used for flow cytometry analysis is provided in Supplemental Figure 2a.

### Toxilight

In selected experiments, Toxilight assays were performed on the supernatant of the functional assays using the Toxilight kit (Lonza) according to manufacturer’s instructions. Wells of effector and target cells alone were included as controls. Target cells were treated with 0.1% Triton-X for 5 minutes at room temperature and supernatant was collected and stored at -20°C. Toxilight assays were run using Toxilight kit (Lonza) on a BioTek Synergy H1 plate-reader (Agilent). Percentage killing was calculated using the following formula:

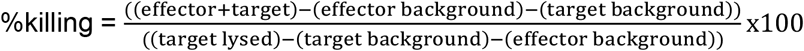

### xCELLigence

Target cells were plated and allowed to sediment and reach a normalised cell index of 1 before the addition of effector cells at a 5:1 ratio 24 hours later. Wells of target cells alone were included as negative controls. Target cells were treated with 0.1% Triton-X as positive controls. Assays were run in triplicates for a total of 120 hours. xCELLigence assays were run on a xCELLigence RTCA MP instrument (Agilent).

### HepG2-NTCP HBV *de-novo* infection

HepG2-NTCP cells were seeded on collagen-coated plasticware. Cells were infected with HBV at a multiplicity of infection (MOI) of 400 genome equivalents per cell in the presence of 4% polyethylene glycol 8,000 (PEG 8,000) for 24 hours. After infection, the viral inoculum was removed, and cells were washed three times with PBS before being maintained in Dulbecco’s Modified Eagle Medium (DMEM) supplemented with 10% FBS. Secreted HBsAg levels were measured by ELISA (Autobio, China), provided in Supplemental Figure 2b.

### Co-culture with T cells

Negative isolation of autologous T cells was performed using CD4 or CD8 isolation kits (Miltenyi Biotec) as per manufacturer instructions using MACS buffer, MidiMACS separator, LS columns (Miltenyi Biotec) and pre-separation filters. Untouched T cells were resuspended in complete RPMI, counted and plated in 96-well U-bottom plates. Half of the wells were stimulated with anti-CD3/anti-CD28 Dynabeads (Gibco, by ThermoFisher Scientific) at a 1:1 ratio for 60 hours.

Cryopreserved expanded NK cells were thawed the day before the co-culture and rested overnight in SCGM media supplemented with 10% human AB serum, 1% L-Glutamine and 100 IU/ml recombinant human IL-2 at 37°C. Negative isolation of NK cells from thawed PBMCs was performed as per manufacturer instructions (NK isolation kit, Miltenyi Biotec) and untouched NK cells were rested in complete RPMI overnight at 37°C.

T cells were washed and co-cultured with expanded NK cells or isolated *ex vivo* NK cells at a 5:1 ratio for 6 hours, in the presence of CD107a and Golgiplug/Golgistop, as described for the functional assays. Cells were stained with fluorochrome conjugated antibodies (listed in Supplemental Table S1) for 20 minutes at 4°C. Cells were washed with PBS before being fixed for 20 minutes at 4°C with Cytofix (Becton Dickenson, Wokingham, UK). Samples were acquired using BD Fortessa x20 using BD FACSDiva 8.0 or Cytek Aurora, using Cytek SpectroFlo software and analysed on FlowJo10.10.0. The gating strategy used for flow cytometry analysis is provided in Supplemental Figure 3a.

### Statistical analysis

Prism 10 (GraphPad) was used for statistical analysis. The Wilcoxon-paired t test was used to compare 2 paired groups. The Friedman test was used to compare 3 paired groups. The statistical significance is indicated in the figures (*p < 0.05, **p < 0.01, ***p < 0.001, and ****p < 0.0001).

## Results

### Expanded NK cells retain adaptive phenotype

We utilised an expansion protocol leveraging K562 HLA-E feeder cells in combination with IL-2, to drive expansion of NKG2C^+^ adaptive NK cells, previously used to expand NK cells from healthy donors [14]. All selected donors with chronic viral infection had high baseline frequencies of NKG2C^+^ NK cells (mean 31%, range 11.7%-57.7%). Their CD3/CD19 depleted PBMCs were co-cultured for 11 days with K562-HLA-E feeder cells pulsed with HLA-G leader peptide (Figure 1a). By day 5, CD14^+^ monocytes and feeder cells were no longer present, and on day 11, the expanded NK cells achieved >97% purity (Figure 1b, Supplemental Figure 1a). The NK cells expanded with a mean of 50-fold, and the NKG2C^+^ proportion of NK cells expanded with a mean of 100-fold during the 11-day culture (Figure 1c). Expanded NK cells displayed a predominant expression of adaptive NK cell markers, with a significant increase in NKG2C expression, compared to the inhibitory counterpart NKG2A (Figure 1d). These populations were characterised by a high expression of inhibitory KIRs (iKIRs), downregulation of the maturation marker CD57 and of the inhibitory receptor Siglec-7 (Figure 1d). Expanded NK cells displayed a significant increase in the expression of activating receptors CD69, DNAM-1, NKG2D and NKp30, and maintained high expression of NKp46. The high levels of Granzyme B expression indicated robust cytotoxic potential. Importantly, expanded NK cells retained CD16 expression and upregulated CD2 expression, both important for ADCC [20] (Figure 1d). These data are in keeping with observations from healthy donors [14]. There was an upregulation of the inhibitory receptor LAG3, with no significant increase in the levels of expression of PD-1, TRAIL or PDL1 (Figure 1e). Due to their phenotypic characteristics typical of adaptive NK cells, expanded NK cells are hereafter referred to as expanded adaptive NK cells (aNK).

**Figure 1.**
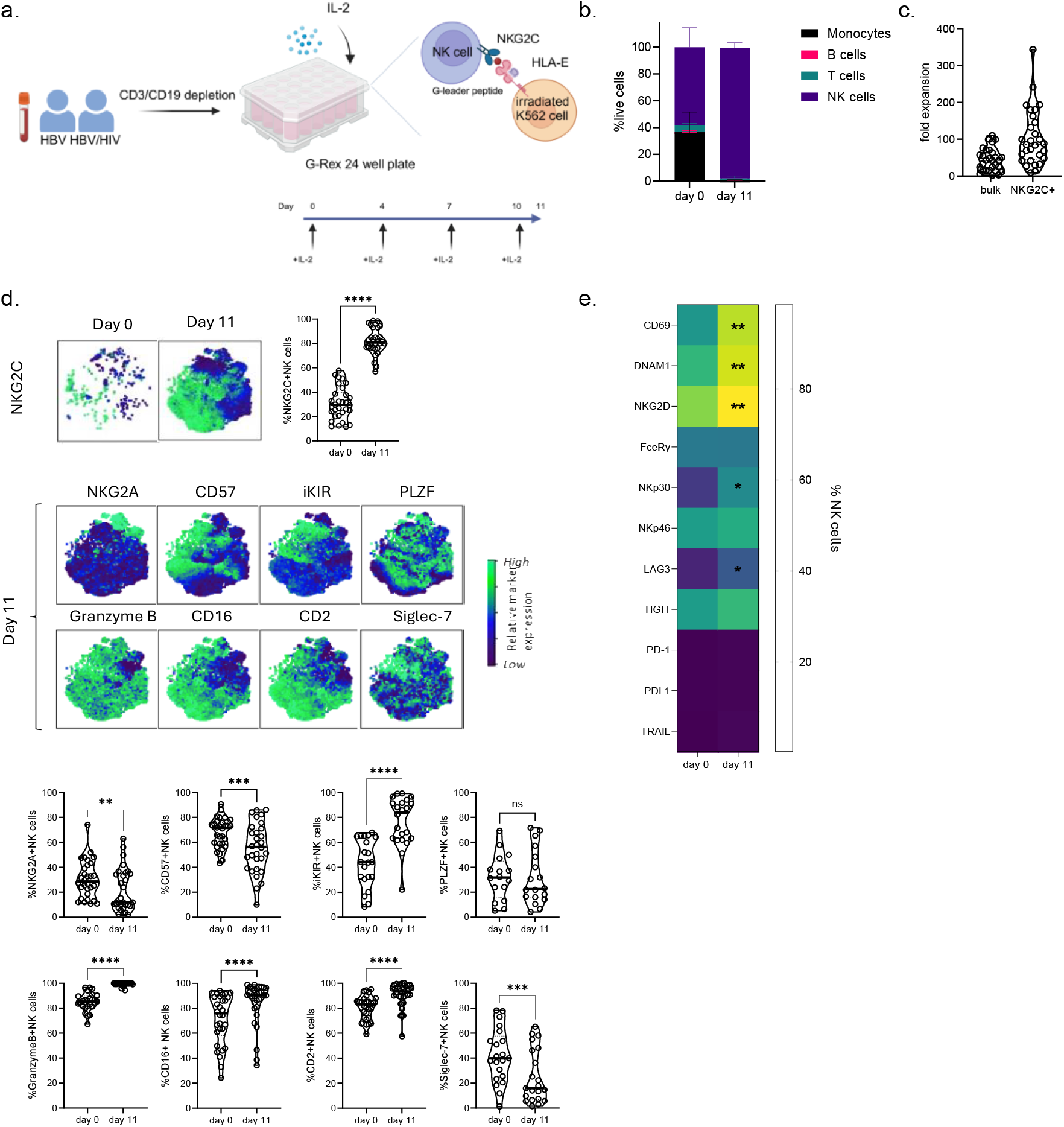
Expanded NK cells display an adaptive phenotype. (a) Expansion protocol; (b) Cell type frequency on day 0 depleted PBMC and day 11 expanded NK cells; (c) Fold expansion in number of total NK cells and NKG2C^+^ NK cells following the 11-day culture; (d) Relative expression of key adaptive markers shown on viSNE clustering and violin plots of sampled events to show frequencies of positive populations; (e) Heatmap of expression levels of depicted NK cell markers on day 0 and day 11. Significance determined by Wilcoxon paired test, *p < 0.05, **p < 0.01, ***p < 0.001 and ****p < 0.0001.

### Expanded aNK cells demonstrate enhanced ADCC capacity

To determine how the phenotypic characteristics of expanded aNK cells translated into functional capacity, we first assessed functional capacity of day 0 vs day 11 expanded aNK cells from the same donors against K562 HLA-E cells. As expected, aNK cells demonstrated enhanced IFNγ expression, degranulation (as measured by the surrogate marker CD107a) and killing (measured via Toxilight assay) against K562 HLA-E cells (Figure 2a and 2b). We then assessed ADCC capacity against Raji cells coated with anti-CD20 antibody. Expanded aNK cells displayed significantly enhanced IFNγ expression and degranulation, as well as killing compared to day 0 counterparts (Figure 2c and 2d). Our results demonstrated increased ADCC activity from expanded aNK cells, in some instances recovering baseline defects.

**Figure 2.**
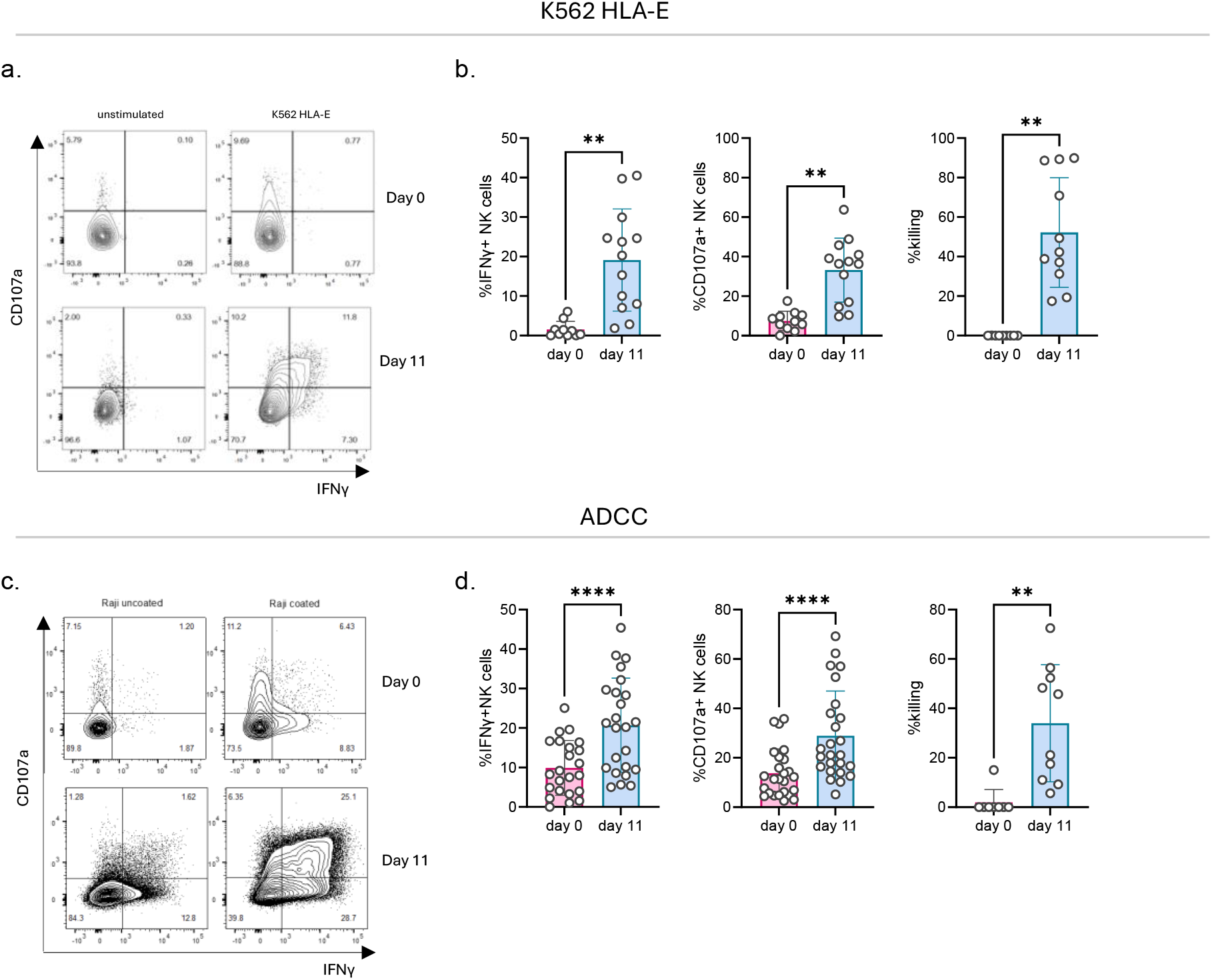
Expanded adaptive NK cells exhibit enhanced ADCC capacity compared to pre-expansion. (a) Representative flow plots of CD107a/IFNy co-expression on bulk NK cells on day 0 and day 11 post-expansion against K562 cells expressing HLA-E; (b) Relative expression of IFNy, CD107a and percentage killing on NK cells against K562 cells expressing HLA-E; (c) Representative flow plots of CD107a/IFNy co-expression on NK cells on day 0 and day 11 post-expansion against Raji cells in the presence of anti-CD20 antibody; (d) Relative expression of IFNy, CD107a and percentage killing of NK cells against Raji cells in the presence of anti-CD20 antibody. Significance determined by Wilcoxon paired test, *p < 0.05, **p < 0.01, ***p < 0.001 and ****p < 0.0001.

### Expanded aNK cells demonstrate enhanced reactivity against hepatoma cell lines and HBV-infected target cells

To explore the potential antiviral capacity of expanded aNK cells, we initially utilised PLC/PRF/5 cells, a hepatoma cell line containing integrated HBV DNA that constitutively expresses and secretes high levels of hepatitis B surface antigen (HBsAg), a known suppressor of NK cell function and cytokine production [21]. This cell line, normally insensitive to NK cell mediated killing, served as our initial model of an HBV-infection *in vitro*. Day 11 expanded aNK cells displayed enhanced degranulation and killing against PLC/PRF/5 cells compared to their day 0 counterparts (Figure 3a). Next, we sought to determine the functional capacity of expanded aNK cells against HepG2-NTCP cells, a hepatoma cell line that normally elicits low-level NK cell responses and expresses the NTCP receptor that HBV uses to gain entry into hepatocytes and thus can be infected with HBV. First, we assessed the reactivity of NK cells against uninfected HepG2-NTCP cells. Our results showed enhanced degranulation and killing against HepG2-NTCP cells relative to day 0 NK cells from the same donors (Figure 3b). Utilising xCELLigence, a real-time cell analysis assay, we continuously monitored target cell growth after addition of effector cells. While the HepG2-NTCP cells continued to grow in the absence of expanded aNK cells, their growth was effectively controlled by the expanded aNK cells, corroborating our findings (Figure 3c).

**Figure 3.**
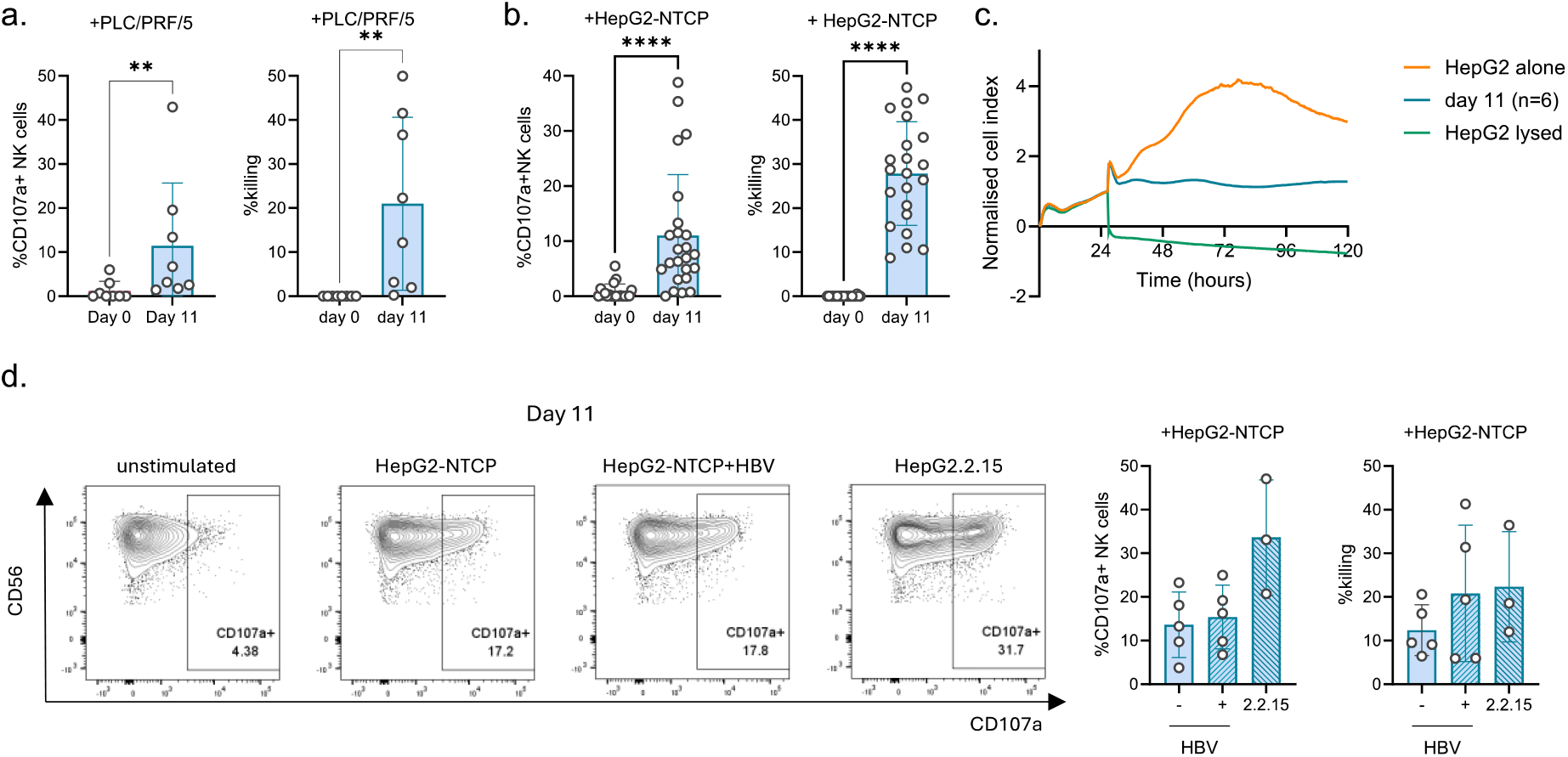
Expanded aNK cells exhibit enhanced functional capacity against HBV-infected HepG2-NTCP cells compared to pre-expansion. (a) Relative expression of CD107a and target cell killing of NK cells against PLC/PRF/5 cells of sampled events; (b) Relative expression of CD107a and target cell killing of NK cells against HepG2-NTCP cells of sampled events; (c) Normalised cell index of HepG2-NTCP cells in the absence or presence of expanded aNK cells; (d) Representative flow plots of CD107a/IFNy co-expression on day 11 expanded NK cells against uninfected HepG2-NTCP cells, HBV-infected HepG2-NTCP cells and HepG2.2.15 cells and relative expression of CD107a and target cell killing against uninfected HepG2-NTCP cells, HBV-infected HepG2-NTCP cells and HepG2.2.15 cells. Significance determined by Wilcoxon paired test, *p < 0.05, **p < 0.01, ***p < 0.001 and ****p < 0.0001.

Finally, we assessed the activity of expanded aNK cells against *de novo* HBV-infected HepG2-NTCP cells and HepG2.2.15 cells, a HepG2-derived cell line stably transfected with replicative HBV DNA that continuously produces viral particles and secretes HBsAg. Secreted HBsAg levels by the infected cell lines are provided in Supplemental Figure 2b. Due to the low responses consistently observed from *ex vivo* NK cells against HepG2-NTCP cells, we focused on the activity of expanded aNK cells. The reactivity of expanded aNK cells against HBV infected HepG2-NTCP cells was comparable to that against uninfected HepG2-NTCP cells, with a trend towards higher degranulation against HepG2.2.15 cells (Figure 3d).

### TGF-β conditioning confers a tissue-resident phenotype and further enhances functional responses in expanded aNK cells

TGF-β is the most abundant immunosuppressive cytokine in the liver microenvironment, contributing to hampered immune cell responses, viral spread and tumour growth [22]. We incorporated low-dose TGF-β in addition to IL-2 in our expansion platform to induce a tissue-resident phenotype and to pre-condition the expanded aNK cells to withstand the immunosuppressive effect of TGF-β encountered in tissue (Figure 4a) [23].

**Figure 4.**
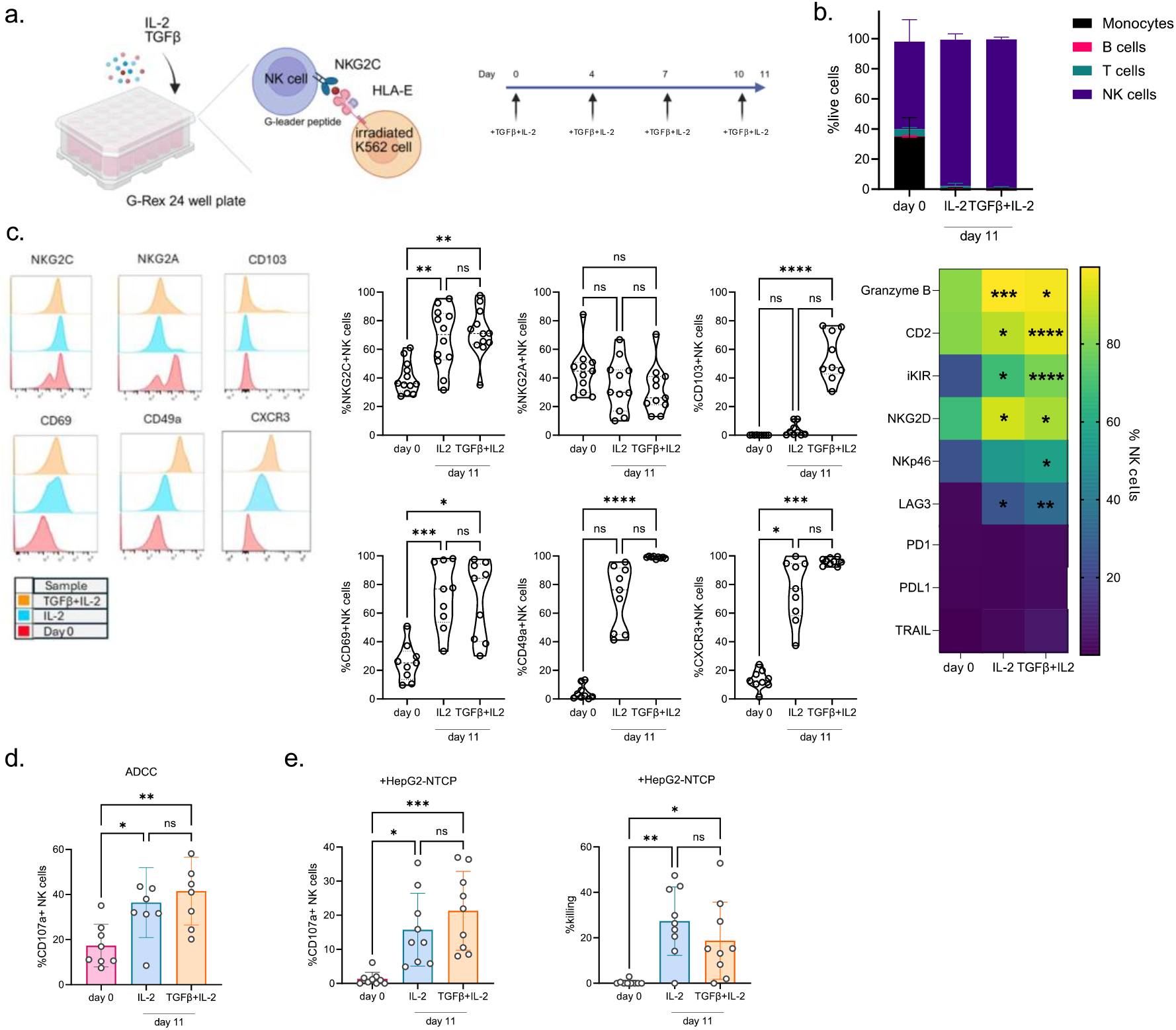
Pre-conditioning with TGF-β results in an expanded aNK cell population with retained adaptive features and adopted tissue-resident phenotype, while maintaining functional responses. (a) Expansion timeline; (b) Cell type frequencies on day 0 depleted PBMC, day 11 IL-2 and day 11 TGF-β+IL-2 expanded aNK cells; (c) Histograms of expression of NKG2C, NKG2A, CD103, CD69, CD49a and CXCR3, summary graphs of expression for these phenotypic markers and heatmap of expression of other phenotypic markers; (d) Summary plot of relative CD107a expression against anti-CD20 coated Raji cells; (e) Summary plot of relative CD107a expression and estimated target cell killing against HepG2-NTCP cells. Significance determined by Friedman paired test, *p < 0.05, **p < 0.01, ***p < 0.001 and ****p < 0.0001. Significance on the heatmap in (c) is shown for comparisons between IL-2 vs day 0 and TGFβ+IL-2 vs day 0.

TGF-β conditioning did not impair the generation of high frequencies of NK cells on day 11 (Figure 4b) and expanded cells displayed a similar adaptive phenotype (high expression of NKG2C, CD2, NKG2D, iKIR and Granzyme B) as IL-2 expanded aNK (Figure 4c). Additionally, these TGF-β imprinted cells adopted high expression of typical NK cell tissue-resident markers CD103, CD49a and CXCR3 (Figure 4c). Importantly, conditioning with TGF-β did not negatively impact ADCC capacity, with degranulation responses remaining comparable to IL-2 expanded aNK cells (Figure 4d). Similarly, responses against HepG2-NTCP cells were enhanced in comparison to day 0 counterparts and were not affected by TGF-β incorporation compared to IL-2 expanded aNK cells (Figure 4e).

### Expanded aNK cells show minimal reactivity against autologous activated T cells

NK cells can target activated immune cells as part of their role in maintaining homeostasis. We and others have demonstrated the capacity of NK cells to kill activated T cells in the liver through various pathways [7, 9]. To evaluate the potential rheostat role of expanded aNK, we co-cultured these cells with autologous anti-CD3/CD28 activated CD4^+^ T cells and measured degranulation (Figure 5a). Expanded aNK cells demonstrated minimal degranulation against autologous activated CD4^+^ T cells compared to *ex vivo* NK cells from the same donors (Figure 5b and 5c). Within the *ex vivo* NK cell pool, the NKG2C^-^ population was responsible for the degranulation observed against activated T cells, while the NKG2C-dominant expanded aNK pool showed no reactivity (Figure 5d and 5e). Similar observations were made against CD8^+^ T cells (Supplemental Figure 3b).

**Figure 5.**
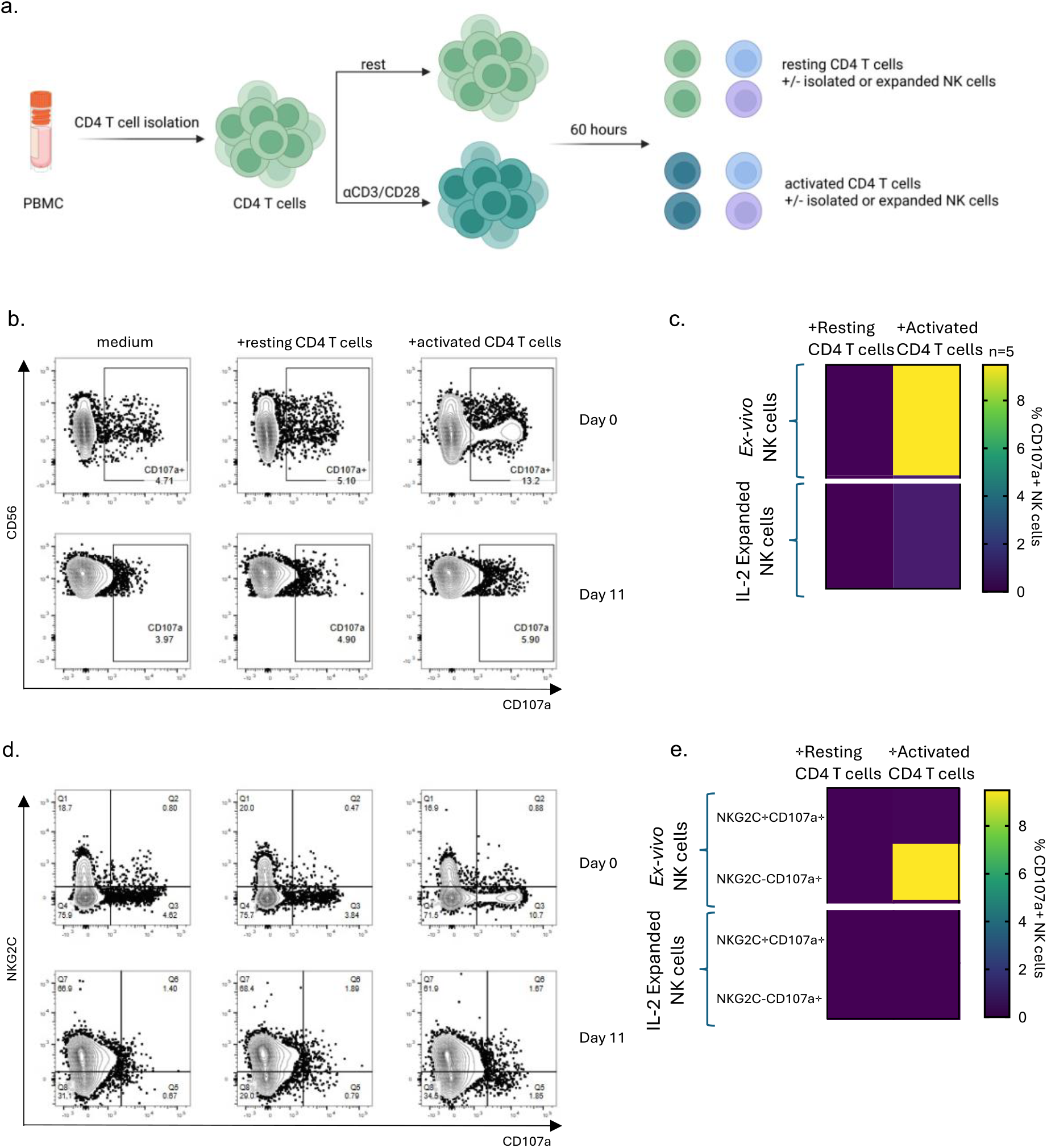
Expanded aNK cells display minimal reactivity to activated CD4^+^ T cells. (a) Assay design; (b) Representative flow plots of CD107a expression of day 0 NK cells and day 11 cryopreserved expanded aNK cells against autologous resting and activated CD4^+^ T cells; (c) Heatmap of CD107a expression on day 0 NK cells and day 11 cryopreserved expanded adaptive NK cells against autologous resting and activated CD4^+^ T cells from 5 donors; (d) Representative flow plots of NKG2C/CD107a co-expression of day 0 isolated *ex vivo* NK cells and day 11 cryopreserved expanded adaptive NK cells against autologous resting and activated CD4^+^ T cells; (e) Heatmap of expression of CD107a according to NKG2C expression of day 0 NK cells and day 11 cryopreserved expanded aNK cells against autologous resting and activated CD4^+^ T cells from 5 donors.

## Discussion

In this study we assessed the robustness of a feeder-based method to enhance both the expansion and functionality of aNK cells in people with chronic viral infection. Our findings provide evidence that such an approach can effectively direct and boost aNK cell subpopulations for functional HBV cure strategies and related HCC.

The expanded aNK cells were characterised by a homogenous receptor expression and high levels of activating receptors (NKG2D, DNAM-1 and CD2) conferring higher sensitivity to target cells. Despite the upregulation of LAG-3 after 11 days in culture, consistent with previous reports [14], the expanded cells retained robust functional capacity, suggesting that this marker may represent activation rather than true exhaustion of expanded aNK cells.

A distinctive advantage of expanded aNK cells, in contrast to other memory-like NK cell populations, such as cytokine induced memory-like (CIML) NK cells, was their lack of TRAIL and PDL1 expression [24], even following TFG-β conditioning (Figure 4c). This is particularly significant in the context of HBV infection, as TRAIL-expressing NK cells have been shown to target both hepatocytes, inducing liver damage, and HBV-specific CD8^+^ T cells, hampering the T cell arm of immunity against HBV [7, 8]. Similarly, PDL1-expressing NK cells can limit therapeutic HBV-vaccine induced CD8^+^ T cell responses [9]. The absence of these potentially detrimental markers on our expanded aNK cells could therefore be advantageous for HBV control by limiting excessive negative regulation of other antiviral effectors and preventing excessive liver damage.

Importantly, our data indicate that expanded aNK cells have a limited regulatory role against autologous activated T cells. This observation aligns with recent findings demonstrating that aNK cells have a reduced capacity for immunoregulatory killing of activated T cells [18]. This characteristic would be particularly beneficial in the context of HBV infection, where robust virus-specific T cell responses are crucial for viral control and ultimate clearance.

TGF-β conditioning resulted in an expanded aNK pool with a tissue-resident phenotype and retained enhanced functional responses. This contrasts with other NK expansion protocols incorporating TGF-β imprinting, where downregulation of activating receptors and upregulation of TRAIL have been observed [25]. The acquisition of tissue-resident markers (CD103, CD49a) and high CXCR3 expression is particularly important for targeting liver-tropic diseases like HBV and HCC, as these markers facilitate tissue homing and retention. Previous studies have shown that NK cells with tissue-resident phenotypes display enhanced anti-tumour activity in solid malignancies [26], suggesting that our TGF-β-conditioned aNK cells may be particularly effective against HCC.

Our findings of enhanced functional responses, robust ADCC activity, and recognition of HCC cell lines in the presence or absence of HBV infection underscore the potential utility of these populations in combination with monoclonal antibodies currently in use as part of functional cure studies. The clinical significance of ADCC in HBV infection has been highlighted by studies showing that antibody-mediated ADCC can target HBV-infected hepatocytes expressing viral antigens [27, 28]. Our expanded aNK cells, with their preserved CD16 expression and enhanced ADCC capacity, could therefore synergise with therapeutic antibodies targeting HBV surface proteins to eliminate infected hepatocytes.

In recent years, NK cells have been recognised for their prognostic and predictive roles across various cancer types, and adoptive NK cell immunotherapy has demonstrated promising results in both pre-clinical and clinical studies targeting solid tumours, including HCC [29, 30]. Recent studies showing that NK cell frequency and function correlate with better outcomes in HCC patients [31, 32], further support the potential benefit of enhanced NK cell responses through adoptive transfer. Although the role of aNK cells specifically in controlling HCC has remained largely unexplored, our data using patient-derived NK cells present a compelling opportunity for autologous therapeutic strategies. Future studies comparing the activity of healthy donor expanded NK cells will further guide the optimal approach for clinical translation.

Other preconditioning techniques to allow NK cells to survive within the tough immunological landscape of the liver will be crucial to maximise the efficacy of expanded NK cells in clinical applications. To that end, we are advancing our expansion platform by incorporating insights from our recent work on mitochondrial dynamics and metabolic reprogramming [33]. By addressing the mitochondrial defects commonly observed in exhausted immune cells within the liver, we aim to enhance the metabolic fitness of aNK cells. This metabolic optimisation approach, combined with their intrinsically elevated bioenergetic profile, may provide aNK cells with the resilience needed for sustained antitumour and antiviral activity in the liver.

As we move forward, additional efforts to improve the expansion platform will be focused on generating a clinical-grade yield of expanded aNK cells. The exquisite specificity of aNK cells to HLA-E presented peptides, which has been shown to influence proliferation and ADCC responses [13], opens the possibility for further optimising these approaches by tailoring the peptide ligand on HLA-E to enhance expansion and selectivity.

Our study has several limitations that should be addressed in future research. First, while we demonstrated enhanced functional responses against hepatoma cell lines, assessment against primary hepatocytes would provide more physiologically relevant insights. Second, *in vivo* studies would be valuable to evaluate the efficacy and safety of expanded aNK cells in relevant animal models of HBV infection. Third, while we observed enhanced functional responses against hepatoma cell lines, the exact mechanisms underlying these improvements require further investigation. Our study concentrated on HCMV-associated aNK cells readily accessible in peripheral blood from our donors; future research will aim to extend these approaches to other unique NK cell subpopulations with liver adaptive/memory features.

In conclusion, our findings provide evidence for the potential role of expanded aNK cells in combating chronic HBV infection and associated malignancies. Our results suggest that leveraging the unique properties of aNK cells could not only improve antiviral and antitumour responses but also provide a novel avenue for HBV functional cure strategies. Future studies will focus on assessing aNK cells against primary hepatocytes and in animal models to facilitate translation into early-phase clinical trials, potentially offering new hope for patients with chronic HBV infection and HBV-related HCC.

## Author Contributions

JK performed experiments, acquisition of data, analysis, and drafting of the manuscript; HAB, BH, AP, NH, KdC performed experiments and contributed to data acquisition and analysis; JD, SM, TF contributed to sample processing; SK, FMB, PK, DM, MKM, USG, AXZ, MWL, KJM, ES contributed to data interpretation and critical editing of the manuscript; PK, UG, MKM, DP recruited donors and obtained blood samples; DP contributed to the conception and design of the study, data interpretation, critical revision of the manuscript, and study supervision.

## Acknowledgements

We thank the participating donors and the laboratory team for their support and contributions.

