## Supplemental Material for "Expanded adaptive NKG2C+ NK cells exhibit potent ADCC and functional responses against HBV-infected hepatoma cell lines"

**Supplemental materials**

**Table S1 –** Flow cytometry antibodies

| Marker | Fluorochrome | Clone | Catalogue number | Supplier |
| --- | --- | --- | --- | --- |
| CXCR5 | BV750 | RF8B2 | 747111 | BD Biosciences |
| CXCR3 | BV650 | G025H7 | 353729 | Biolegend |
| CXCR6 | AF647 | K041E5 | 356008 | Biolegend |
| Lag-3 | eFluor506 | 3D5223H | 69-2239-42 | Invitrogen |
| CD16 | BUV496 | 3G8 | 612944 | BD Biosciences |
| CD49a | BUV563 | SR84 | 749263 | BD Biosciences |
| NKG2D | BUV615 | 1d11 | 751232 | BD Biosciences |
| CD103 | BUV661 | Ber-ACT8 | 749993 | BD Biosciences |
| CD56 | BUV737 | NCAM16.2 | 612766 | BD Biosciences |
| CD8 | BUV805 | SK1 | 612889 | BD Biosciences |
| NKp46 | Pacific Blue | 9E2 | 331912 | Biolegend |
| CD2 | BV480 | S5.2 | 746645 | BD Biosciences |
| CD3 | BV510 | SK7 | 344828 | Biolegend |
| CD57 | BV605 | NK-1 | 567205 | BD Biosciences |
| HLA-DR | BV570 | L243 | 307637 | Biolegend |
| PD-1 | BV421 | eh12.2h7 | 329920 | Biolegend |
| TIM-3 | BV711 | F38-2E2 | 345024 | Biolegend |
| CD14 | Spark Blue550 | 63D3 | 367148 | Biolegend |
| LAIR-1 | PerCP-Cy5.5 | NKTA255 | 342804 | Biolegend |
| KLRG1 | PerCp-eFluor710 | 13F12F2 | 46-9488-42 | eBioscience |
| NKG2C | PE | S19005E | FAB138P-100 | R&D |
| CD4 | cFluor yg584 | SK3 | R7-20041 | Cytek Bioscience |
| TIGIT | BV421 | A15153G | 372716 | Biolegend |
| CD69 | PE-Cy5 | FN50 | 310908 | Biolegend |
| CD159a (NKG2A) | PE-Cy7 | Z199 | B10246 | Beckman Coulter |
| PDL-1 | PE-Fire-810 | 29E.2A3 | 329755 | Biolegend |
| KIR2DL1/S1/S3/S5 | APC | HPMA4 | 339510 | Biolegend |
| KIR3DL2 |  | 539304 | FAB2878A | R&D |
| KIR2DL2 |  |  | A22333 | Beckman Coulter |
| KIR3DL1 |  |  | 130-092-474 | Miltenyi Biotec |
| CD19 | Spark NIR 685 | HIB19 | 302270 | Biolegend |
| CD38 | PE-FIRE 700 | S17015A | 397121 | Biolegend |
| FCeRγ | FITC |  | FCABS400F | Millipore |
| Granzyme B | AF700 | GB11 | 560213 | BD Biosciences |
| NKG2D | FITC | 1d11 | 320820 | Biolegend |
| DNAM-1 | BUV563 | DX11 | 748429 | BD Bioscience |
| NKp30 (CD337) | BV650 | P30-15 | 743171 | BD Bioscience |
| TRAIL | BV421 | RIK-2 | 564243 | BD Biosciences |
| CD57 | Pacific Blue | HNK-1 | 359608 | Biolegend |
| CD107a | BV605 | H4A3 | 328634 | Biolegend |
| TNFα | FITC | MAb11 | 502906 | Biolegend |
| CD19 | BV510 | HIB19 | 740164 | BD Biosciences |
| CD14 | BV510 | M5E2 | 301842 | Biolegend |
| CD4 | BV510 | OKT4 | 317444 | Biolegend |
| CD16 | BB700 | 3G8 | 746199 | BD Biosciences |
| CD56 | BV605 | NCAM16.2 | 562780 | BD Biosciences |
| CD3 | BV650 | OKT3 | 317324 | Biolegend |
| CD2 | APC | RPA-2.10 | 300214 | Biolegend |
| CD57 | PE-Dazzle594 | HNK-1 | 359620 | Biolegend |
| CD107a (LAMP-1) | APC-H7 | H4A3 | 561343 | BD Biosciences |
| IFNγ | BV421 | B27 | 562988 | BD Biosciences |
| TNFα | BV711 | MAb11 | 502940 | Biolegend |
| CD328 (Siglec-7) | APC/Fire 750 | 6-434 | 339207 | Biolegend |
| CD2 | BV711 | RPA-2.10 | 300231 | Biolegend |
| CD7 | PE/Cyanine5 | CD7-6B7 | 343110 | BD Biosciences |
| CD57 | BV421 | NK-1 | 563896 | BD Biosciences |
| PLZF | PE-CF594 | R17-809 | 565738 | BD Biosciences |
| CD38 | BV785 | HIT2 | 303530 | Biolegend |
| Live/Dead | Aqua | N/A | L34957 | Thermo Fisher Scientific |
| Live/Dead | Blue | N/A | L23105 | Thermo Fisher Scientific |

**Supplemental figures**

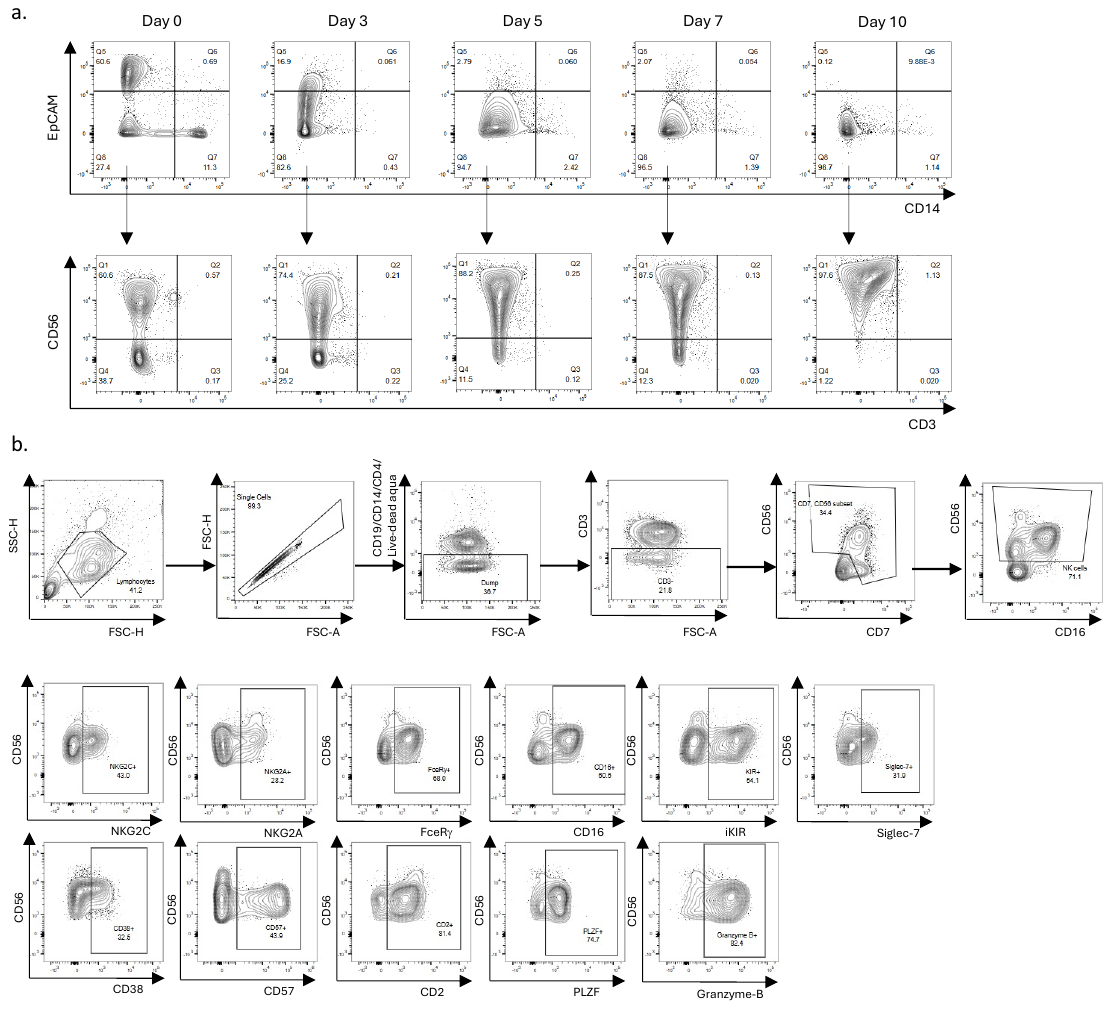

**Supplemental Figure 1.** (a) Culture composition on day 0, 3, 5, 7 and 10; (b) gating strategy for phenotypic analysis incorporated in Figure 1. This gating strategy shows a pre-depletion PBMC sample.

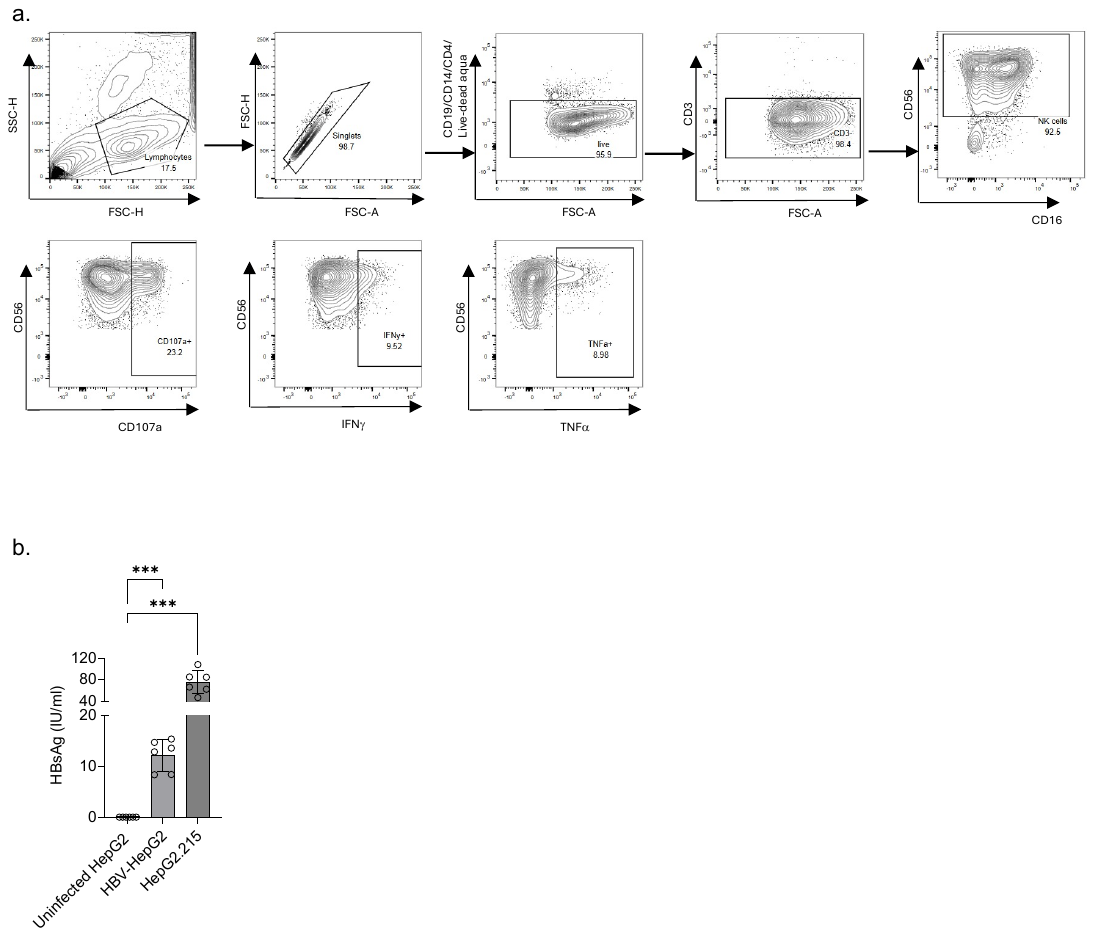

**Supplemental Figure 2.** (a). Gating strategy used for functional assays. This gating strategy shows expanded aNK cells against HepG2-NTCP; (b). HepG2-NTCP cells were infected with HBV at a multiplicity of infection (MOI) of 400, or HepG2.215 cells (an HBV integrated cell line) were cultured for 6 days. Supernatants were collected and analysed for HBsAg levels using an ELISA assay (mean ± SEM, n = 6, One-way ANOVA with multiple comparisons, Two-sided).

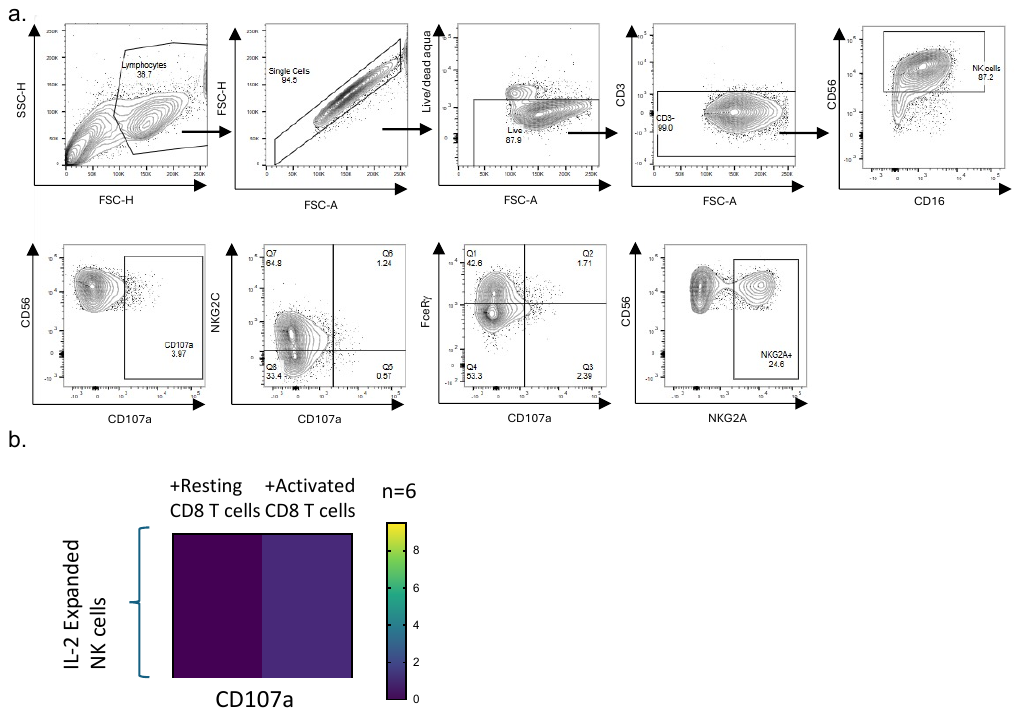

**Supplemental Figure 3.** (a) Gating strategy used for NK-T cell co-culture experiments in Figure 5; (b) Heatmap of expression of CD107a of day 0 isolated *ex vivo* NK cells and day 11 expanded adaptive NK cells against autologous resting and activated CD8^+^ T cells from 6 donors.
